# Partial-resistance against aphids in wild barley involves phloem and mesophyll-based defences

**DOI:** 10.1101/502476

**Authors:** Daniel J Leybourne, Tracy A Valentine, Jean AH Robertson, Estefania Pérez-Fernández, Angela M Main, Alison J Karley, Jorunn IB Bos

## Abstract

Aphids, including the bird cherry-oat aphid (*Rhopalosiphum padi*), are significant agricultural pests. Aphid populations are typically controlled using insecticides, but there is increasing demand for more sustainable pest management practices. The wild relative of barley, *Hordeum spontaneum* 5 (Hsp5) has been described as partially-resistant to *R. padi*. Partial-resistance is proposed to involve higher thionin and lipoxygenase gene expression. However, the underlying mechanistic processes are unknown. In this study we compared Hsp5 with a susceptible cultivar of barley (Concerto) to test the extent to which partial-resistance affects aphid fitness. We used the electrical penetration graph technique to monitor *R. padi* feeding patterns to elucidate the tissue location of partial-resistance factors alongside molecular and biochemical analyses to identify potential mechanisms. We show that partial-resistance in Hsp5 extends to three aphid species and is mediated by phloem/mesophyll-based factors, leading to a three-fold increase in the time aphids take to establish sustained phloem ingestion. Partial-resistance likely involves elevated expression of defence and phytohormone genes alongside altered phloem amino acid composition. Further work is required to establish the function of these traits, however this study highlights plant tissues which are important in conferring broad-spectrum partial-resistance against aphids in barley.

**Highlight:** Partial-resistance against aphids in wild barley is based in the mesophyll and vascular tissue and is potentially associated with higher basal defence gene expression and altered phloem amino acid composition.

## Introduction

Barley, *Hordeum vulgare*, is the fourth most agriculturally important cereal by production quantity (Newton et al., 2011), with 65% of barley yield being used as feed for livestock, 30% in malting and distilling processes, and 5% contributing towards other uses (Newman and Newman, 2006). By 2050 it is estimated that crop yields will need to increase by 60% in order to meet the demands of global population growth (Alexandratos and Bruinsma, 2012). One way to help achieve this is to reduce yield losses to abiotic and biotic stress factors. Aphids such as the bird cherry-oat aphid, *Rhopalosiphum padi*, and the English grain aphid, *Sitobion avenae*, are major pests of cereals. Aphids feed directly from plant phloem using specialised mouthparts known as stylets (Auclair, 1963). Aphids cause direct plant damage by feeding and ingesting phloem sap and act as vectors of economically important plant viruses. Many cereal aphids are vectors for viruses such as *Barley Yellow Dwarf Virus (BYDV*), which are transmitted into plant tissues during stylet probing (Powell, 2005). Disease resulting from virus infection can cause yield losses of up to 30% in barley and up to 80% in wheat (Smith and Sward, 1982; Perry et al., 2000; Murray and Brennan, 2010).

Aphids are primarily controlled through the application of insecticides, with the amount of insecticide used per annum increasing (Jess et al., 2018). However the use of these has led to the emergence of insecticide-resistant aphid populations (Chen et al., 2007; Foster et al., 2014), caused negative impacts on non-target organism (Unal and Jepson, 1991; James et al., 2016), and potentially resulted in an increase in *BYDV* prevalence (Dewar and Foster, 2017), leading to legislative restrictions on insecticide use and promotion of integrated pest management (Directive, 2009/128/EC). As a consequence, there is a growing requirement for alternative pest management solutions which are more sustainable, and improving crop resistance to aphids is one avenue which could be explored to achieve this.

To date, only partial-resistance against aphids has been reported for cereal crops. Plant resistance traits which are effective against insect pests can be allocated to one of three categories: 1) chemical deterrence of insect settling and feeding; 2) physical barriers to insect feeding; and 3) reduction in palatability (Mitchell et al., 2016). Partial-resistance against aphids has been associated with each of these categories (respectively, Gibson and Pickett (1983), Tsumuki et al. (1989) and Greenslade et al. (2016). Inherent molecular defences also play a key role in conferring resistance against aphids (Delp et al., 2009; Guan et al., 2015; Escudero-Martinez et al., 2017). Molecular defences shown to be involved in aphid resistance in *Hordeum* spp. include increased expression of thionin (antimicrobial peptides) genes (Delp et al., 2009; Mehrabi et al., 2014; Escudero-Martinez et al., 2017), increased chitinase and ß-1,3-glucanase activity (Forslund et al., 2000), and the presence of plant secondary metabolites (Gianoli and Niemeyer, 1998). Plant phythormone signalling pathways, including Abscisic Acid (ABA), Salicylic Acid (SA), Jasmonic Acid (JA) and Ethylene (ET) signalling processes, mediate co-ordinated molecular responses to herbivory via the regulation of defence signalling genes and the biosynthesis of defensive allelochemicals (Smith and Boyko, 2007; Bari and Jones, 2009; Morkunas et al., 2011; Foyer et al., 2016); higher constitutive expression of phytohormone signalling genes can lead to improved resistance against aphids in cereals (Losvik et al., 2017).

Constitutive and induced plant defences are frequently lost during plant breeding and domestication (Moreira et al., 2018). However, wild relatives of modern crops possess traits which confer improved resistance against herbivorous pests (Dempewolf et al., 2014). Screening crop wild relatives for (partial-)resistance against aphids provides an opportunity to identify potentially beneficial traits (Zhang et al., 2017) which can be introduced into agricultural cultivars to improve resistance (Xu et al., 2015; Li et al., 2018). Screening of wild relatives in wheat (Xu et al., 2015; Aradottir et al., 2017), barley (Nieto-Lopez and Blake, 1994; Åhman et al., 2000) and maize (Maag et al., 2015) has resulted in successful identification of partial-resistance traits in wild relatives of these species (Tsumuki et al., 1989; Barria et al., 1992; Gianoli and Niemeyer, 1998; Ahmad et al., 2011; Greenslade et al., 2016; Chandrasekhar et al., 2018; Li et al., 2018).

Partial-resistance against *R. padi* was identified in the barley wild relative *H. spontaneum* 5 (Hsp5) (Åhman et al., 2000; Delp et al., 2009). Although Hsp5 features higher tissue concentrations of indole alkaloid gramine, a plant defensive compound, the presence of gramine did not correlate with increased partial-resistance (Åhman et al., 2000). Instead, partial-resistance in Hsp5 is thought to involve increased expression of plant defence genes (Delp et al., 2009; Mehrabi et al., 2014), including higher constitutive expression of thionin genes (Delp et al., 2009) and higher expression of a proteinase inhibitor (Mehrabi et al., 2014). Initial attempts have been made to characterise the molecular processes contributing to partial-resistance in Hsp5 (Delp et al., 2009), although the underlying mechanism(s) of resistance and the tissue location of these resistance factors remains to be elucidated.

The primary aim of this study was to characterise the plant traits and mechanisms contributing to partial-resistance in Hsp5 by comparison with a susceptible commercial barley cultivar, Concerto, and explore whether this partial-resistance is effective against several aphid species, including two agricultural pests (*R. padi* and *Sitobion avenae*) and an invasive species (*Utamphorophora humboldti)*. The factors contributing towards partial-resistance in Hsp5, and their tissue location, were elucidated using the electrical penetration graph (EPG) technique to monitor *R. padi* feeding processes. Subsequently, we assessed differences between the two barley species in leaf surface architecture, defence gene expression, and phloem amino acid composition to gain insight into the potential mechanisms underlying partial resistance in Hsp5.

## Materials and methods

### Plant growth and insect rearing conditions

*Hordeum vulgare* Linnaeus cv. Concerto (Concerto) and *H. spontaneum* 5 Linnaeus (Hsp5) seeds were surface-sterilised by washing in 2% (v/v) hypochlorite and rinsing with d.H_2_O. Seeds were kept moist in the dark: Hsp5 seeds were incubated at 4°C for 14 days and Concerto seeds were kept at room temperature for 48h. Germinated seedlings were planted into a bulrush compost mix (Bulrush, Northern Ireland) and grown under glasshouse conditions with a 16:8h day:night length and a 20:15°C day:night temperature. Plants were grown until the first true leaf stage (1.1 – 1.2 on the Zadoks et al. (1974) growth staging key).

Asexual laboratory cultures of the bird cherry-oat aphid, *Rhopalosiphum padi* (Linnaeus) (genotype B; Leybourne et al. (2018)), the English grain aphid, *Sitobion avenae* Fabricius and the American grass leaf aphid, *Utamphorophora humboldti* Essig were established from individual apterous adults collected from Dundee, UK. Molecular barcoding of the cytochrome oxidase subunit I gene (Folmer et al., 1994) was used to confirm identity of aphid species. Aphid cultures were reared on one week old barley seedlings (cv. Optic) contained in ventilated cups at 20°C and 16:8h (L:D).

### Insect fitness measurements

The performance of all aphid species was assessed in glasshouse conditions (described above) on Hsp5 and Concerto (*n* = 12) in a randomised block design (each block contained one replicate of each treatment combination). Plants were infested with a single apterous aphid which was allowed to reproduce overnight, then a total of three nymphs were retained on each plant. Mean nymph mass and number of surviving nymphs were recorded at 72h and 168h, after which a random single nymph was returned to the plant; for this nymph, data were collected on the length of the pre-reproductive period (d) and the intrinsic rate of population increase (r_m_). Aphids were caged onto the first fully expanded true leaf using Perspex clip-cages of 25mm internal diameter (MacGillivray and Anderson, 1957). Nymph mass gain was calculated as the change in mass between 168 and 72h and aphid r_m_ was calculated using the equation of Wyatt and White (1977): 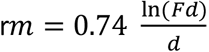, where d is the time period between birth and production of first progeny, and Fd is the total progeny produced over a time period equal to d.

### Electrical penetration graph (EPG) monitoring of aphid feeding

The DC-EPG technique (Tjallingii, 1978; Tjallingii, 1988; Tjallingii, 1991; Tjallingii, 2001) was employed to monitor the probing and feeding behaviour of apterous *R. padi* over a 6h period using a Giga-4 DC-EPG device (EPG Systems, The Netherlands) on plants at the true-leaf stage. A plant probe (copper rod approximately 50 mm long x 5 mm diameter), was soldered to electrical wire extending from the plant voltage output of the Giga-4 device and inserted into the plant soil. An aphid probe was made by soldering a piece of copper wire (30 mm long x 2 mm diameter) to a brass pin (tip diameter 2 mm). Approximately 30 mm of gold wire (20 μm diameter; EPG Systems, The Netherlands) was adhered to the copper end of the aphid probe using water-based silver glue (EPG Systems, The Netherlands) and aphids were connected by adhering the free end of the gold wire onto the aphid dorsum using the same water-based adhesive. Wired aphids were connected to the Giga-4 device by placing the end of the brass pin into the EPG probes with a 1 GΩ input resistance and a 50x gain (Tjallingii, 1988). The order in which *R. padi* – plant combinations were tested and allocated to an EPG probe was randomised. Data were acquired using Stylet+D software (EPG Systems, The Netherlands). A total of 18 and 16 successful recordings were made for aphids feeding on Concerto and Hsp5, respectively. All EPG recordings were obtained within a grounded Faraday cage.

EPG waveforms were annotated using Stylet+A software (EPG Systems, The Netherlands). Waveforms were annotated by assigning waveforms to np (non-probing), C (stylet penetration/pathway), pd (intercellular punctures), E1 (saliva secretion into phloem), E2 (saliva secretion and passive phloem ingestion), F (penetration difficulty) or G (xylem ingestion) phases (Tjallingii, 1988; Alvarez et al., 2006). No E1e (extracellular saliva secretion) phases were detected. Annotated waveforms were converted into time-series data using the excel macro developed by Dr Schliephake (Julius Kühn-Institut, Germany).

### Determination of non-glandular trichome density

Non-glandular trichome densities of Hsp5 and Concerto (*n* = 9) were determined using polarised light microscopy following a procedure adapted from Pomeranz et al. (2013). Briefly, the first true leaf was excised from plant stems and treated to two incubation steps to clear the leaf epidermis. Firstly, leaf area was measured and leaves were soaked in 96% Ethanol (Sigma-Aldrich, UK) for 48h before being treated with 1.25 M NaOH:EtOh (1:1, v:v) at 70°C for 2h. Leaves were stored in 50% glycerol prior to trichome visualisation; if NaOH:EtOH treatment did not clear the leaf epidermis leaves were further treated with 80% lactic acid for 6h. To analyse trichome density samples were placed adaxial side up on microscope slides (75 x 25 mm; Corning, UK). A polarising light microscope was created using two 50 mm^2^ polarising filters (Sigma-Aldrich, UK), one placed below the stage of a stereo-microscope but above the light source and the second attached below the objective lens. Non-glandular trichomes appeared illuminated under the polarised microscope setup and the number of trichomes per unit area were counted. To allow correlation of trichome density with aphid performance, an additional *R. padi* performance experiment (as described above) was carried out on these plants prior to trichome density analysis.

### Epicuticular wax analysis

A Fourier Transform Infrared (FTIR) spectrometer (Bruker Vertex 70 FTIR spectrometer, Bruker, Ettlingen, Germany), incorporating a Diamond Attenuated Total Reflection (DATR) sampling accessory, was used to identify the functional groups present in chemical extracts from the surfaces of Hsp5 and Concerto leaves. Briefly, the waxes from upper and lower leaf surfaces of the first true leaf (n = 4) were extracted with dichloromethane (DCM) by running approximately 1-2 ml of DCM along the leaf surface directly onto the diamond window of the ATR sampling accessory; after evaporation of the DCM, FTIR spectra were recorded of the films deposited on the DATR. Signal-to-noise ratio was enhanced by taking 200 scans for each sample and averaging to obtain a single spectrum. The spectral range scanned was 4000 cm^-1^ to 400 cm^-1^ and a background reading was taken before each sample was analysed.

### Analysis of phloem amino acid composition

Hsp5 and Concerto plants were grown under glasshouse conditions (as described above) in a temporally-split randomised block design: three temporal blocks, one initial temporal block comprising two replicate sub-blocks, and two further temporal blocks each comprising four replicate sub-blocks. Each sub-block contained a single replicate of every treatment combination. Plants at the first true-leaf stage were infested with either ten apterous *R. padi* adults caged onto the plants with microperforated bags (Polybags, UK) or were aphid-free bagged controls. Samples were collected at 0 and 24h post-aphid infestation. Phloem sap was collected using the method of King and Zeevaart (1974); leaves (*n* = *10*) were excised from the stem, placed in a dish containing 1 mM EDTA (Sigma-Aldrich, UK) solution and immediately re-cut. The cut surface of the leaf was placed into 200μl of filter-sterilised 1mM EDTA solution (pH 7.5 with NaOH) in a 2 mL Eppendorf™ microcentrifuge tube and incubated for 1.5h at room temperature in a darkened exudation chamber (a polystyrene box containing a dish of 6 M K_2_HPO_4_ (Sigma-Aldrich, UK), and moistened paper towels to maximise humidity and decrease leaf transpiration). After incubation, leaves were removed and the samples of EDTA solution containing exuded phloem sap were stored at −80°C until analysis.

Reverse-phase HPLC was used to separate phloem amino acids Asp, Glu, Asn, His, Ser, Gln, Arg, Gly, Thr, Tyr, Ala, Trp, Met, Val, Phe, Ile, Leu and Lys using an Agilent HP1100 series autosampling LC system equipped with a ZORBAX™ Eclipse AAA column and fluorescence detector. Amino acids were derivatised using *o*-phthaldialdehyde (Sigma-Aldrich, UK) following the method developed by Jones et al. (1981); all protein amino acids except proline and cysteine could be detected by this method, with a detection limit of approximately 0.5 pmol. Samples were processed in their experimental blocks along with blanks (1 mM EDTA solution that was not exposed to an excised leaf) as controls. Amino acids were identified and quantified by comparison with known concentrations of amino acid standards (AA-S-18 (Sigma, UK), supplemented with Asp, Glu and Trp).

### qPCR analysis of plant defence genes

Five sub-blocks over the three temporal blocks (one from the first block and two from each subsequent temporal block) were chosen at random from the phloem exudation experiment described above and leaf tissue was sampled for gene expression analysis. Approximately 25mm length of leaf tissue was cut from the true leaf and flash-frozen in liquid nitrogen immediately prior to the excision of the leaf for phloem exudation. Plant material was stored at −80°C until RNA extraction.

For RNA extraction leaf samples were ground to a fine powder under liquid nitrogen using a mortar and pestle and total RNA was extracted with the Norgen Plant/Fungi RNA Extraction Kit (Sigma-Aldrich, UK) following the manufacturer’s protocol with additional DNAse I treatment (Qiagen, UK); RNA quality and quantity was assessed with a NanoDrop^®^ ND-1000 Spectrophotometer (ThermoFischer, UK). cDNA synthesis was carried out with approximately 1000 ng RNA using the SuperScript^®^ III cDNA synthesis kit (Sigma-Aldrich, UK), following the manufacturer’s protocol. qPCR primers were designed using the Roche Universal Probe Library Assay Design Centre. Reference gene primers were described by Hua et al. (2015). All primer sequences are shown in Supplementary Table 1 and were supplied by Sigma-Aldrich, UK. Primers were validated for PCR efficiency prior to use, and reference gene primers were tested for stability across all treatments following the geNorm procedure (Vandesompele et al., 2002). Gene expression analysis used SYBR^®^ Green chemistry with GoTaq^®^ qPCR Master Mix (Promega, UK) on a StepOne™ Real -Time PCR Machine (Applied Biosystems, UK). Reactions were carried out in 12.5 μl reactions with a final concentration of 1x GoTaq^®^ qPCR Master Mix, 1μM of each primer, 1.4 mM MgCl_2_, 2.4 μM CXR reference dye and a cDNA quantity of approx. 12.5 ng (assuming 1:1 RNA:cDNA conversion). The qPCR conditions were as follows: 95°C for 15 mins followed by 40 cycles of denaturing for 15s at 95°C, annealing for 30s at 60°C, and 30s at 72°C for DNA extension. Fluorescence was recorded at the end of each annealing cycle and a melting curve was incorporated into the end of the qPCR programme. Data were normalised to the mean expression of two reference genes, *HvCYP* (AK253120.1) and *HvUBC* (AK248472.1), and 2^-ΔΔCt^ methodology (Livak and Schmittgen, 2001) was used to determine differential expression, with Concerto at 0h aphid infestation used as the experimental control treatment. Expression levels at 24h were further normalised to uninfested control plants. Samples were processed in experimental blocks with three technical replicates at the PCR level.

### Statistical analysis

All statistical analyses were carried out using R Studio v.1.0.143 running R v.3.4.3 (R Core Team, 2014) using packages: car v.2.1-4 (Fox and Weisberg, 2011), coxme v.2.2-7 (Therneau, 2018), dunn.test v.1.3.5 (Dinno, 2017), ggplot2 v.2.2.1 (Wickham, 2009), ggpubr v. 0.1.2 (Kassambara, 2017), ggfortify v.0.4.5 (Tang et al., 2016), lawstat v.3.2 (Hui et al., 2008), lme4 v.1.1-13 (Bates et al., 2015), lmerTest v.2.0-33 (Kuznetsova et al., 2017), lsmeans v.2.27-62 (Lenth, 2016), survival v.2.41-3 (Therneau and Grambsch, 2000), survminer v.0.4.2 (Kassambara and Kosinski, 2017), vegan v.2.5-3 (Oksanen et al., 2013).

Aphid juvenile mass gain and adult r_m_, were modelled using linear mixed effects models with experimental block incorporated as a random factor. ANOVA with type III Satterthwaite approximation for degrees of freedom was used to analyse the final models with calculation of Least Squares Means used for *post-hoc* testing. A Cox proportional hazards regression model was used for nymph survival analysis (Therneau and Grambsch, 2000; Therneau, 2018), incorporating experimental block as a random factor; a χ^2^ test was used on the final model. A log-rank test with Benjamini-Hochberg correction was used to carry out pairwise comparisons between aphid-plant combinations. Aphid feeding behaviour was assessed globally by fitting a permutated MANOVA to the dataset. Response variables with normal data distribution were analysed by either ANOVA or general linear models, while non-normally distributed data were analysed with Kruskal-Wallis tests.

Levene’s test (Hui et al., 2008) was used to analyse differences in trichome density; a Kendall’s rank correlation tau was then used to test for correlations between trichome density and *R. padi* performance. Differences in leaf surface chemistry were analysed using a Welch two sample *t*-test by comparing the total number of identified functional groups in the two plant species.

Phloem amino acid concentrations (pmol/μl) were expressed as relative amino acid composition by converting into mole % of total amino acid content and subjected to principal component analysis with a correlation matrix. Three principal components explained 69% of observed variation and were analysed by fitting a linear mixed effects model incorporating aphid treatment, plant, time-point, plant x time-point interaction, and leaf subsampling for RNA (to test effect on amino acid composition) as explanatory variables with temporal block, randomised block and HPLC batch as random factors. Final models were analysed with a χ^2^ test. Differential gene expression (2^-ΔΔct^) data were analysed using Kruskall-Wallis rank-sum tests, with subsequent Dunn’s test *post-hoc* analysis of the plant x infestation interaction.

## Results

### A wild relative of barley exhibits partial-resistance against multiple aphid species

Aphid performance experiments were undertaken to assess whether partial-resistance in Hsp5 against *R. padi* (Delp et al., 2009; Leybourne et al., 2018) extends to other aphid species. We assessed the fitness of *R. padi, S. avenae*, and *U. humboldti* on Hsp5 in comparison with a commercial barley cultivar (Concerto) and found evidence for partial-resistance against all three aphid species. Specifically, nymph survival was significantly lower on Hsp5 (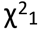 = 10.65; p = 0.001; Fig. 1A; Table 1); pairwise comparison showed that nymph survival was reduced on Hsp5 for *U. humboldti* (p = <0.001), but not for *R. padi* (p = 0.057) or *S. avenae* (p = 0.973). Furthermore, nymph mass gain (F_56,1_ = 9.14, p = 0.003; Fig. 1B; Table 1) and the rate of population increase (r_m_) (F_54, 1_ = 27.43, p = <0.001; Fig. 1C; Table 1) were reduced for all aphid species when feeding from Hsp5. Differences were also detected between aphid species, with *R. padi* exhibiting the highest r_m_ (F_54,2_ = 99.10, p = <0.001; Fig. 1B; Table 1) and *S. avenae* the largest mass gain (F_56,2_ = 49.82, p = <0.001; Fig. 1C; Table 1). On average *U. humboldti* was the least fit species (Fig. 1A-C).

**Fig 1.**
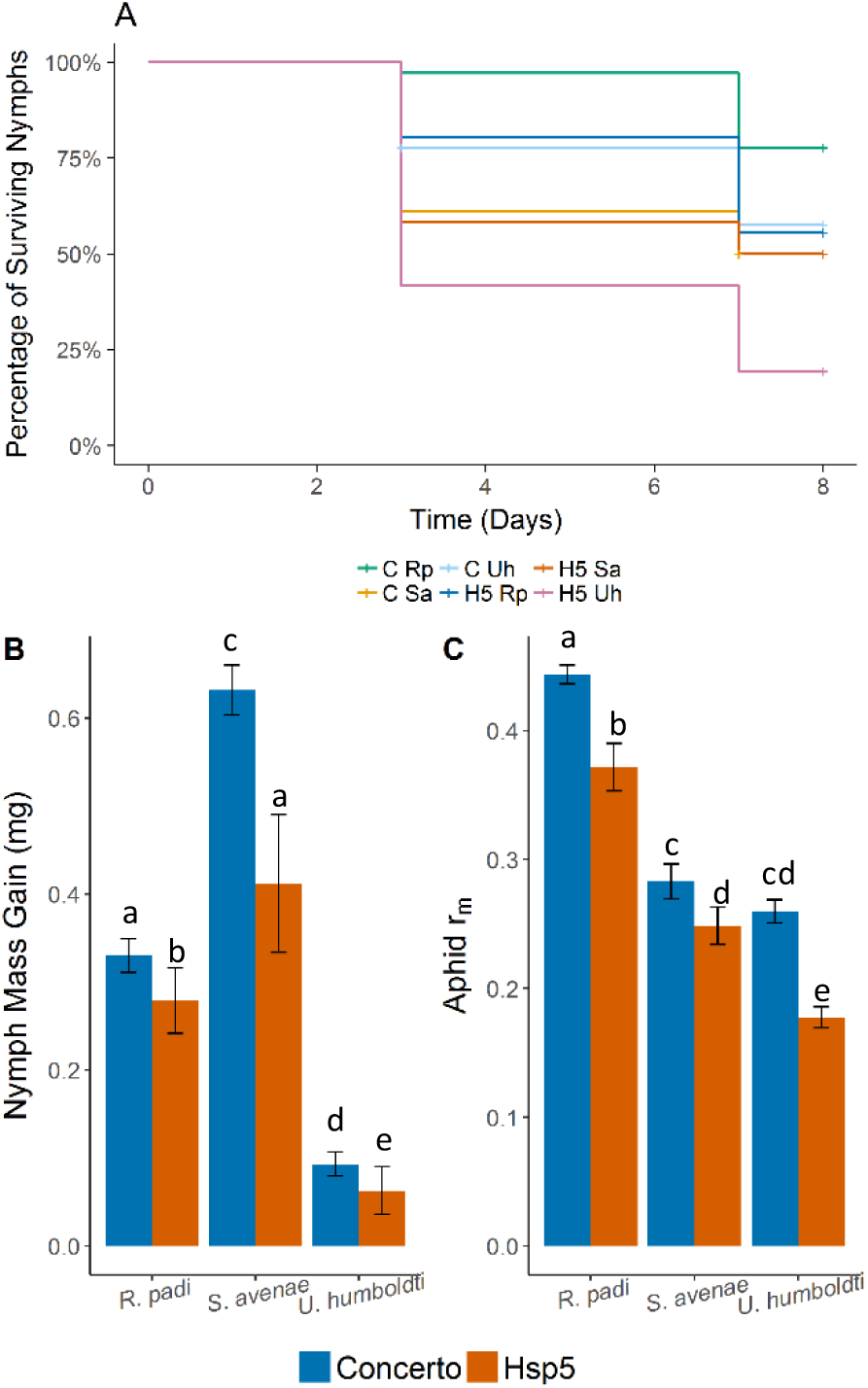
Aphid performance on a susceptible barley cultivar (Concerto) and partially-resistant wild relative (HsP5). A) Survival of aphid nymphs over seven days. Number of model observations = 216. C = Concerto; H5 = Hsp5; Rp = *R. padi;* Sa = *S. avenae;* Uh = *U. humboldti*. B) Nymph mass gain and C) intrinsic rate of population increase (r_m_) of the three aphid species while feeding from the two plant types. Values represent means ± SE. Number of model observations = 72. Letters indicate which groups are similar to each other based on least squares means analysis with Tukey correction.

**Table 1:** Statistical results of aphid performance experiments (significant values are shown in bold).

| Response Variable | Explanatory Variable | Model Basis | Error Distribution or Statistical Method | Statistical Test | Test Statistic | P value |
| --- | --- | --- | --- | --- | --- | --- |
| Nymph Survival | Plant | Cox Proportional Hazards Regression | N/A | Type II Wald $\chi^2$ Analysis of Deviance | $\chi^2_1 = 10.65$ | **0.001** |
| | Aphid | | | | $\chi^2_2 = 13.94$ | **<0.001** |
| | Plant x Aphid | | | | $\chi^2_2 = 5.85$ | 0.054 |
| Nymph mass gain (mg) | Plant | Linear Mixed Effects Model | Restricted Maximum Likelihood | Type III Analysis of Variance with Satterthwaite approximation for degrees of freedom | $F_{1,56} = 9.14$ | **0.004** |
| | Aphid | | | | $F_{2,56} = 49.83$ | **<0.001** |
| | Plant x Aphid | | | | $F_{2,46} = 3.12$ | 0.054 |
| Rate of population increase ($r_m$) | Plant | | | | $F_{1,54} = 27.43$ | **<0.001** |
| | Aphid | | | | $F_{2,54} = 99.10$ | **<0.001** |
| | Plant x Aphid | | | | $F_{2,52} = 1.49$ | 0.234 |

### Partial-resistance against R. padi involves mesophyll and phloem resistance factors

To elucidate the potential underlying partial-resistance mechanism(s) against aphids in Hsp5 *R. padi* feeding behaviour was monitored using the EPG technique. Overall, the feeding behaviour of *R. padi* differed significantly when feeding from Hsp5 compared with Concerto (F_1,33_ = 2.61; p = 0.022; Fig. 2; Table 2). Aphids showed significant differences in feeding behaviour at the leaf epidermis and within mesophyll tissue (Fig. 3; Table 2). We observed an approximate three-fold decrease in the time to the first epidermal probe when feeding on Hsp5 (F_1,33_ = 7.99; p = 0.008), alongside a shorter duration of the first probe (F_1,33_ = 6.94; p = 0.013) and an increased time to the first sieve element puncture (F_1_,_33_= 6.33; p = 0.017). In addition, aphids feeding on Hsp5 showed a two-fold delay in initiating passive phloem ingestion (χ^2^_1,33_ = 4.29; p = 0.038) and the number of intracellular punctures observed during the first aphid probe of plant tissue was lower, with an average of ten intracellular probes, compared with 24 on Concerto (F_1,33_ = 3.95; p = 0.047; Table 2).

**Fig. 2:**
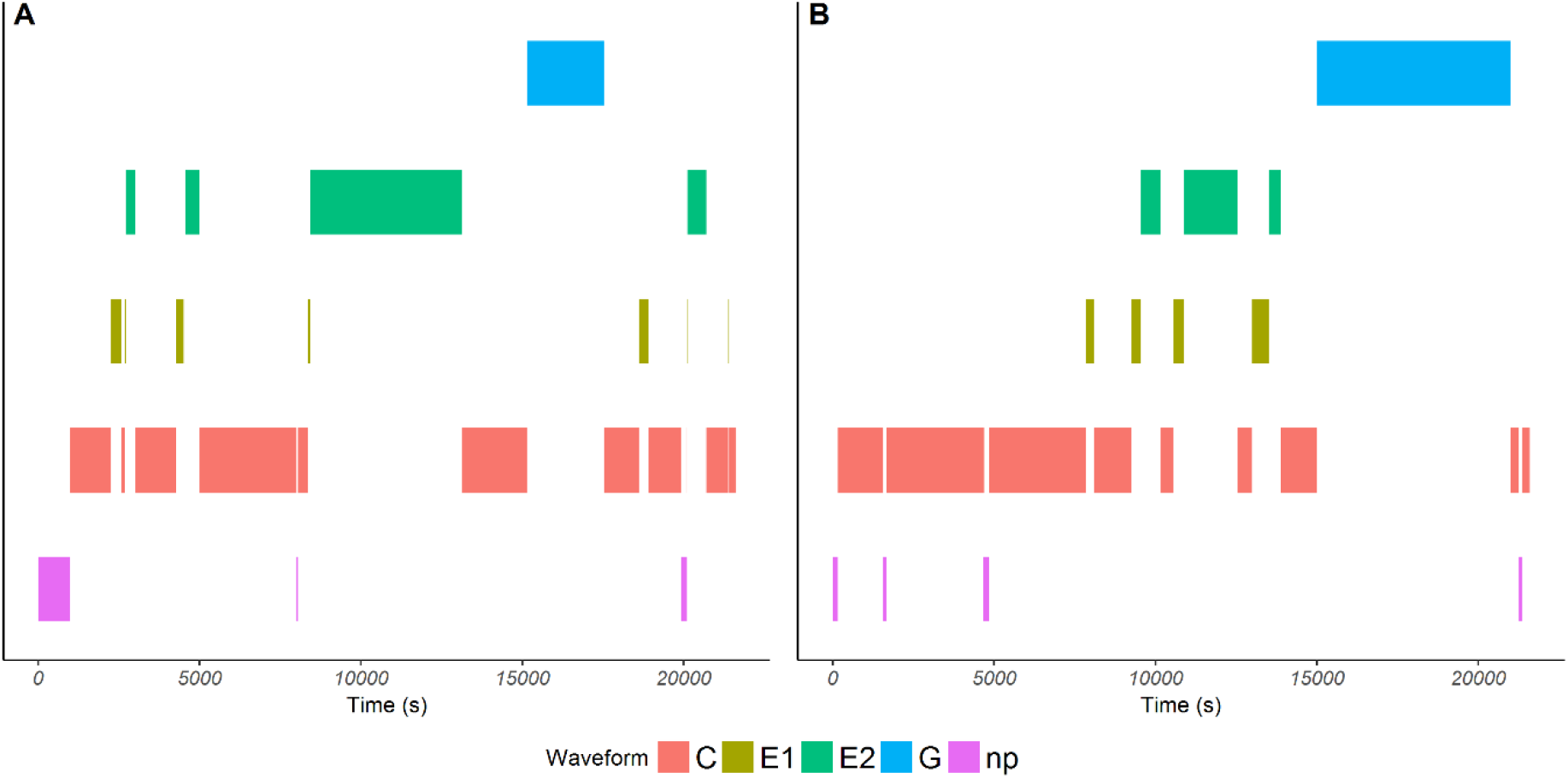
Representative examples of feeding patterns derived from EPG waveforms of *R. padi* feeding from susceptible Concerto (A) and partially-resistant Hsp5 (B). Diagrams show the frequency and length of each feeding parameter over a 6h (21600s) period. Feeding patterns are split into the five main waveforms observed: non-probing of plant tissue (np), probing into epidermal and mesophyll tissue (the pathway phase (C), saliva secretion into phloem (E1), phloem ingestion (E2), and xylem ingestion (G).

**Table 2:** Statistical results of the significant EPG parameters. The mean value for each plant host and the standard error of the mean are also displayed; in the test statistic column χ^2^ values are italicised and F values are underlined.

| EPG Parameter | Hypothesised location of resistance factor | Transformation | Mean Value |  | Standard Error of the Mean |  | Statistical Test | Test Statistic | Degrees of Freedom (Residuals) | P Value |
| --- | --- | --- | --- | --- | --- | --- | --- | --- | --- | --- |
|  |  |  | cv. Concerto | HsP5 | cv. Concerto | HsP5 |  |  |  |  |
| All parameters (Global analysis) | All tissue | - | - | - | - | - | Permutated Multiple Analysis of Variance | <u>2.61</u> | 1(33) | **0.022** |
| Time to the first aphid probe of plant tissue | Epidermis | Log[2] | 1211 s | 421 s | 354 s | 85 s | Analysis of Variance (ANOVA) | <u>7.99</u> | 1(33) | **0.008** |
| Duration of the first aphid probe | Mesophyll | Log[10] | 7717 s | 2147 s | 1737 s | 1161 s | ANOVA | <u>6.94</u> | 1(33) | **0.013** |
| Number of potential drops (intracellular punctures) in first probe | Mesophyll | Sqrt | 24 | 10 | 5.36 | 2.82 | ANOVA on glm | 3.95 | 1(33) | **0.047** |
| Time to the first E1 phase (sieve element puncture and salivation into phloem) | Mesophyll | Log[2] | 2856 s | 7452 s | 660 s | 1683 s | ANOVA | <u>6.33</u> | 1(33) | **0.017** |
| Time to the first phloem phase containing E1 and/or E2 | Mesophyll | - | 3761 s | 9864 s | 675 s | 1884 s | Kruskal-Wallis Test (KW) | 5.50 | 1(33) | **0.019** |
| Time to the first phloem ingestion (E2) phase | Mesophyll | - | 4395 s | 9946 s | 846 s | 1880 s | ANOVA | <u>5.32</u> | 1(33) | **0.028** |
| Time to first sustained phloem ingestion (> 10 minutes) | Mesophyll | - | 8210 s | 13660 s | 1345 s | 1954 s | KW | 4.29 | 1(33) | **0.038** |
| Number of probes before first E1 | Mesophyll | - | 0.66 | 2.38 | 0.34 | 0.76 | KW | 9.36 | 1(33) | **0.002** |
| Number of probes before first E2 | Mesophyll | - | 1 | 2.5 | 0.38 | 0.79 | KW | 4.14 | 1(33) | **0.042** |
| Total time ingesting xylem | Xylem | - | 2197 s | 6859 s | 721 s | 1408 s | KW | 5.28 | 1(33) | **0.022** |
| Total number of xylem phases | Xylem | - | 0.72 | 1.56 | 0.13 | 0.30 | KW | 4.77 | 1(33) | **0.029** |
| Total time ingesting phloem (E2) | Phloem | - | 7965 s | 3299 s | 1306 s | 1188 s | KW | 8.00 | 1(33) | **0.004** |
| Average time of phloem ingestion (E2) | Phloem | - | 5317 s | 1717 s | 1533 s | 702 s | KW | 6.52 | 1(33) | **0.011** |
| Total time of phloem phases containing E1 and/or E2 | Phloem | - | 8824 s | 3620 s | 1348 s | 1241 s | KW | 9.22 | 1(33) | **0.002** |
| Maximum time of a single phloem phase with E1 and/or E2 | Phloem | - | 7913 s | 3225 s | 1442 s | 1172 s | KW | 9.01 | 1(33) | **0.002** |
| Maximum length of a single E2 period | Phloem | - | 7215 s | 2884 s | 1371 s | 1134 s | KW | 8.40 | 1(33) | **0.004** |
| Average time of phloem phase containing E1 and/or E2 | Phloem | - | 5911 s | 1913 s | 1653 s | 722 s | KW | 6.88 | 1(33) | **0.008** |
| Time from first E1 phase to sustained E2 phase | Phloem | - | 8210 s | 13660 s | 1345 s | 1954 s | KW | 4.29 | 1(33) | **0.038** |
| Total number of sustained E2 phases | Phloem | - | 1.39 | 0.75 | 0.16 | 0.17 | KW | 6.06 | 1(33) | **0.014** |
| Mean duration of sustained E2 | Phloem | - | 6500 s | 2779 s | 1424 s | 1141 s | KW | 7.12 | 1(33) | **0.008** |
| Median duration of sustained E2 | Phloem | - | 6425 s | 2779 s | 1440 s | 1141 s | KW | 6.40 | 1(33) | **0.011** |
| Total time of sustained E2 | Phloem | - | 7694 s | 3060 s | 1342 s | 1162 s | KW | 7.49 | 1(33) | **0.006** |
| Ingestion:Pathway Ratio (%) | Phloem | - | 3.01 | 0.67 | 1.40 | 0.33 | KW | 7.43 | 1(33) | **0.006** |

**Fig 3:**
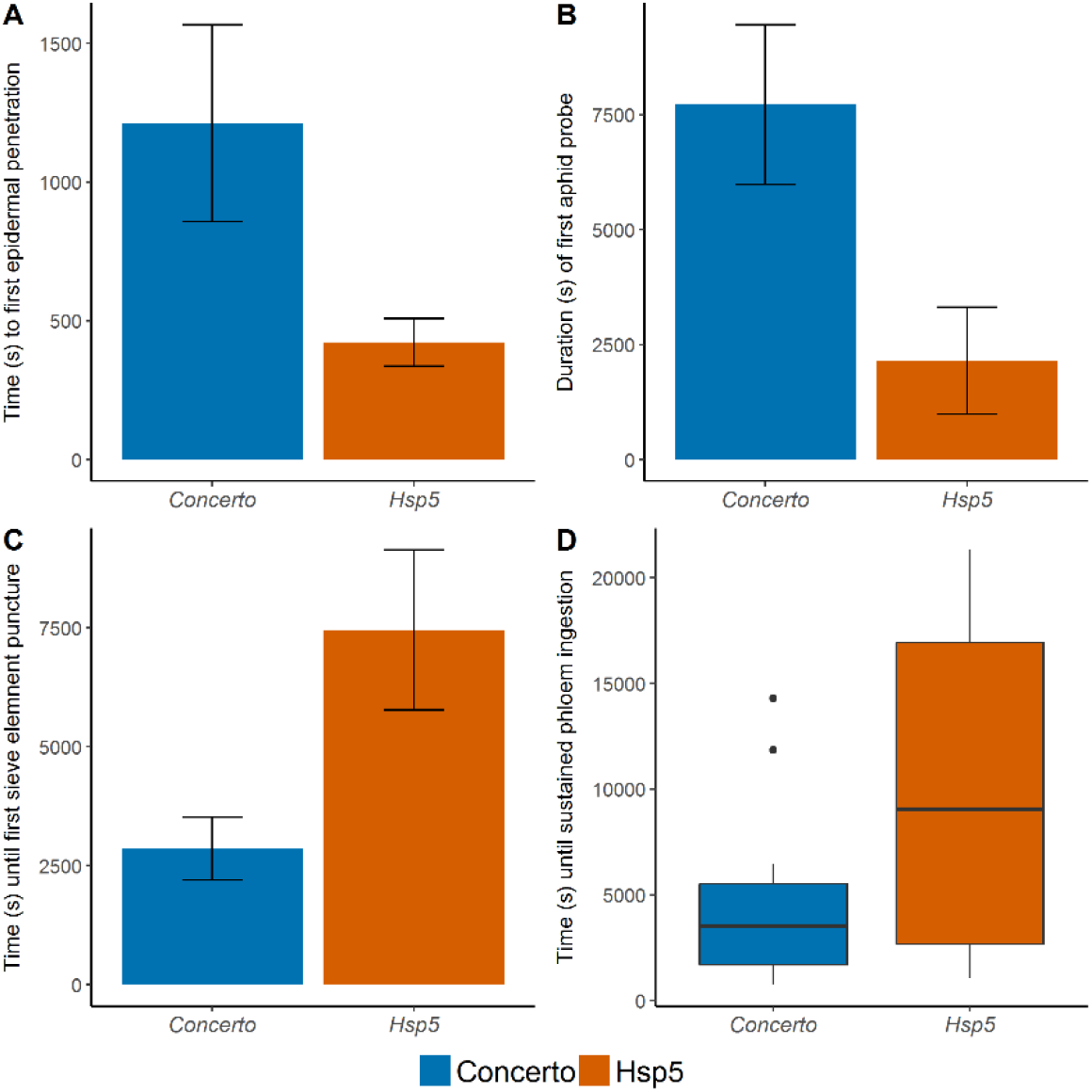
EPG parameters indicative of epidermal, mesophyll and mesophyll/phloem resistance factors. A) Time (s) from start of EPG recording to first stylet puncture of leaf tissue. B) Duration (s) of first stylet probe into plant tissue. C) Time (s) since start of EPG recording until stylet puncture of a sieve tube element. D) Time (s) from start of EPG recording until sustained phloem ingestion. A, B and C values represent means ± SE. Number of model observations = 34.

Feeding patterns within the vascular tissue also differed (Fig. 4; Table 2). Specifically, aphids feeding on Hsp5 showed delayed initiation of sustained phloem ingestion (ingestion for >10 mins) after the first sieve element puncture (χ^2^_1,33_ = 4.29; p = 0.038). The total length of time aphids spent in the sustained feeding phase was also 2.5x shorter (χ^2^_1,33_ = 7.49; p = 0.006), and the ratio of time aphids spent ingesting phloem relative to the length of time probing plant tissue (ingestion:pathway ratio) was four-fold lower (χ^2^_1,33_ = 7.43; p = 0.006). Furthermore, aphids feeding on Hsp5 spent three times longer ingesting xylem (χ^2^_1,33_ = 5.28; p = 0.022). Table 2 reports additional feeding parameters which highlight further mesophyll and vascular-mediated resistance against *R. padi* in Hsp5.

**Fig 4:**
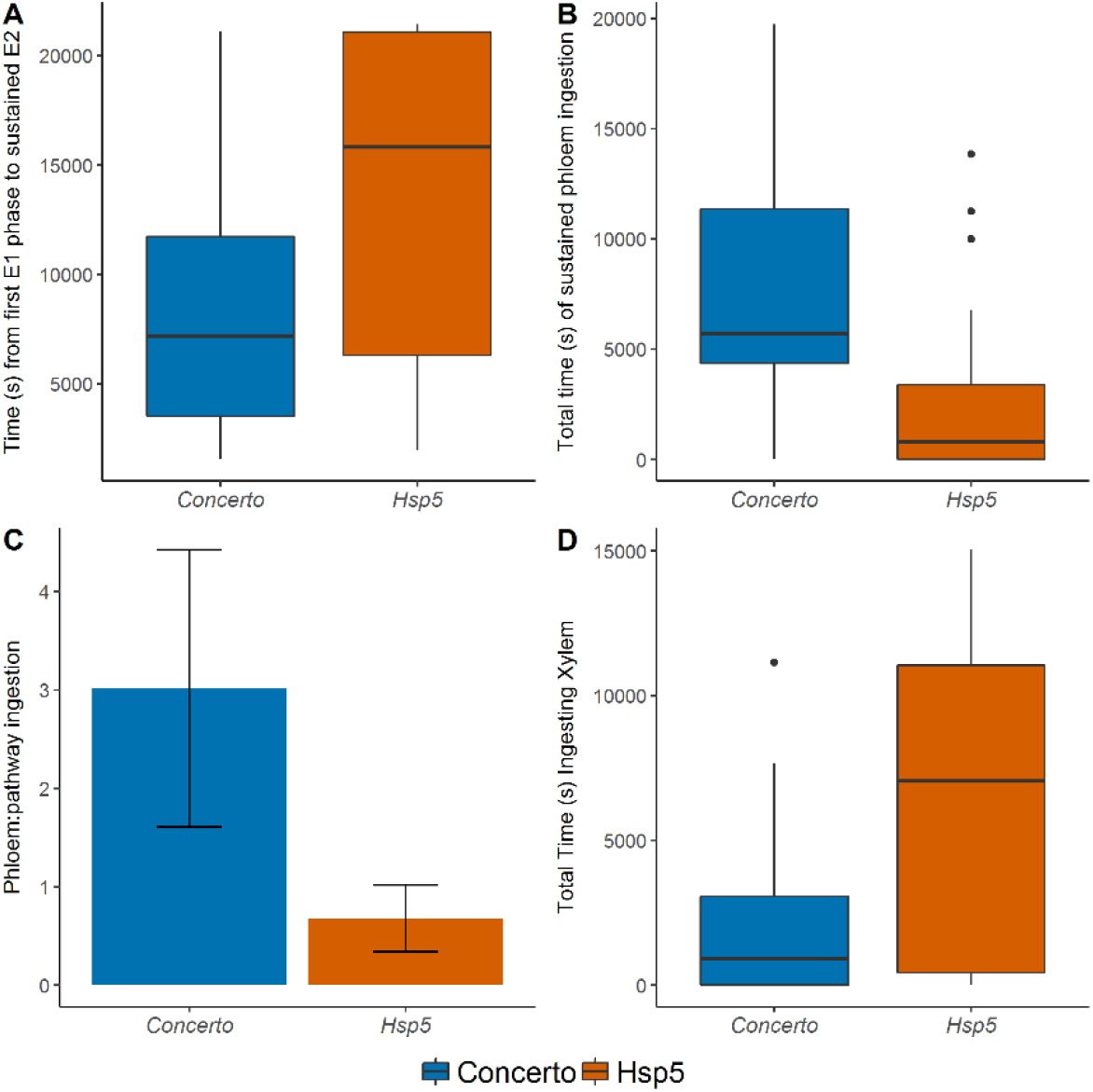
EPG parameters indicative of vascular (phloem and xylem) resistance factors. A) Time (s) from first stylet penetration of a sieve tube element to first sustained phloem ingestion. B) Total time (s) of sustained phloem feeding. C) Ratio of time spent in phloem phase relative to the pathway phase; values represent means ± SE. D) Total time (s) ingesting xylem. Number of model observations = 34.

### Leaf surface architecture differs between susceptible and partially-resistant plants

The leaf surface represents a key interface between plants and insects, with many factors reported to modulate plant-insect interactions, including leaf trichomes and epicuticular waxes (Agrawal et al., 2009; Glas et al., 2012; Karley et al., 2016). Leaf trichome counts revealed Hsp5 had a significantly higher abundance of non-glandular leaf trichomes (F_1,18_ = 6.24; p = 0.022; Fig. 5). Non-glandular trichome abundance showed a negative correlation with *R. padi* fecundity (Z = −3.60; p = <0.001; T = −0.56; Supplementary Fig. 1A) but there was no relation with nymph mass gain (Z = −0.29; p = 0.770; T = −0.04; Supplementary Fig. 1B). In addition, we found differences in the epicuticular wax composition between the two plants. The complexity of the epicuticular wax differed in that aliphatic hydrocarbons were detected in surface extracts of Hsp5 leaves, two Hsp5 replicates also contained weak ester bands (Supplementary Table 3). In Concerto the chemical mixture contained a more consistent mixture of aliphatic hydrocarbons and more pronounced ester bands, alongside a more complex, but variable, mixture of chemical groups, including carboxylates and amides (*t* = – 4.89; df = 6; p = 0.002). Overall, the results were consistent within Hsp5 but more variable in Concerto (Supplementary Table 3). However, the detection of some Nitrogen-containing functional groups in Concerto leaf extracts could be a result of extracting some components within the upper tissue layers, this potentially highlights differences in the thickness of the wax between Hsp5 and Concerto.

**Fig. 5:**
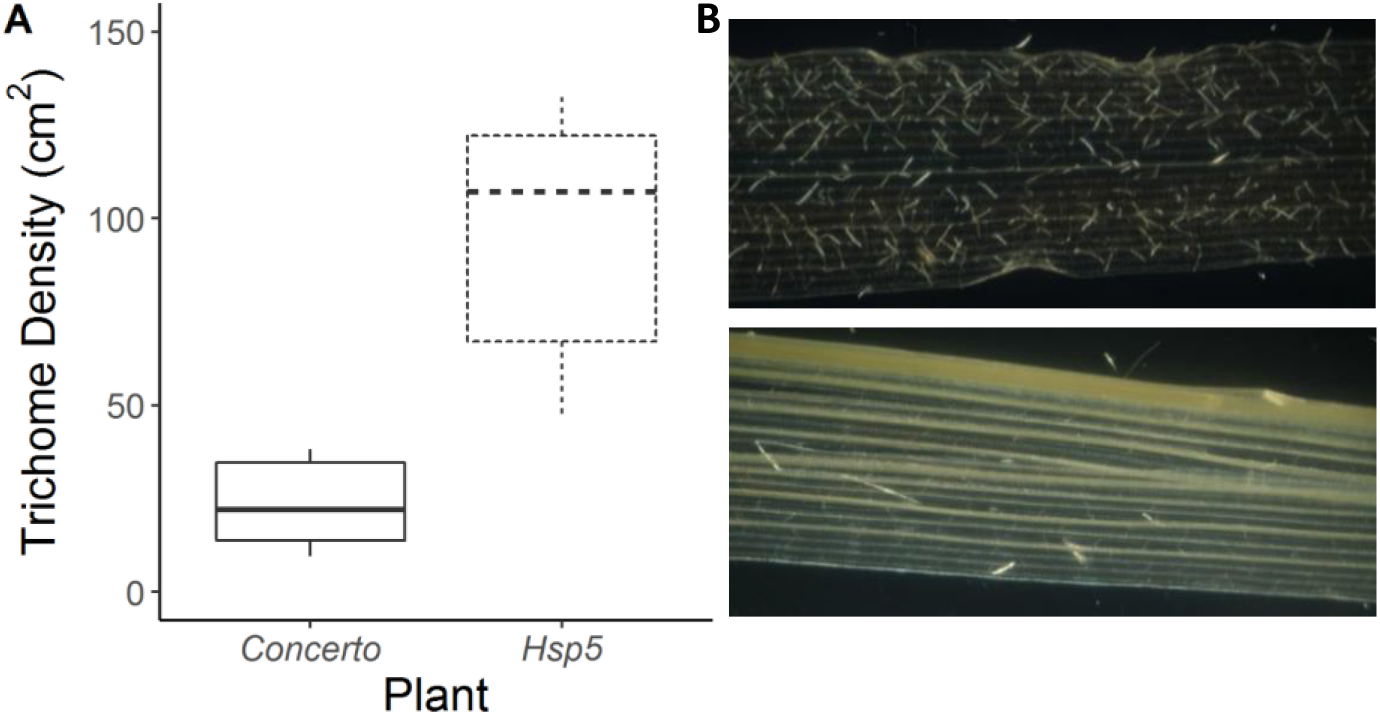
Trichome densities in the two plant types. A) Median non-glandular trichome densities (No. cm^-2^) and confidence intervals for Concerto and Hsp5. B) Polarised light micrograph showing the contrasting leaf hair densities between HsP5 (top) and Concerto (bottom). Number of model observations = 30.

### Basal expression levels of several plant defence genes are elevated in Hsp5

To investigate the level of defence gene expression in Concerto versus Hsp5 we selected marker genes for defensive thionins (*HvTHIO1, HvTHIO2, HvßTHIO*) alongside JA (HvLOXA, *HvLOX2* and *HvJAZ*), SA (*HvNPR1*), ET (*HvERF*), and ABA (*HvAl*) signalling pathways. We assessed expression of marker genes in both plant types constitutively and in response to 24h of aphid infestation. We observed *10-(HvTHIO1), 13-(HvTHIO*) and 7-fold (*HvßTHIO*) higher expression of the three thionin genes in Hsp5 (Fig 6; Table 3). Thionin gene expression levels remained higher in Hsp5 24h after aphid infestation but were not differentially regulated in either plant in response to aphid infestation (Fig 6; Table 3). In addition, *HvLOXA, HvLOX2, HvA1*, and *HvERF1*, were more highly expressed in Hsp5, with 3.5-, 6-, 10– and 14-fold higher expression levels, respectively (Fig 6; Table 3).

**Fig 6.**
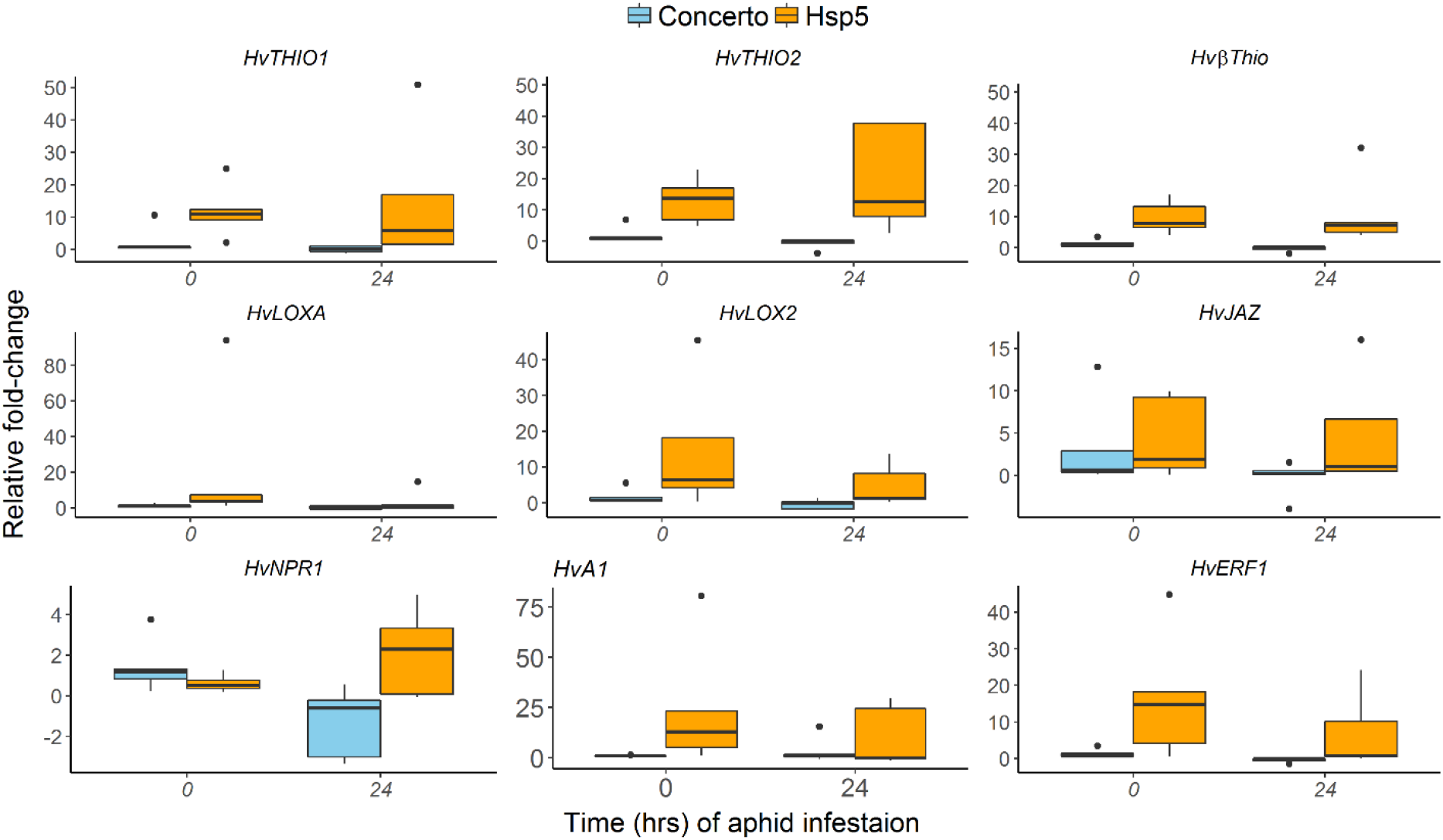
Expression patterns of thionin and phytohormone signalling genes 0 h and 24 h after aphid infestation. All gene expression values are relative to the mean expression of two reference genes, *HvUBC*and *HvCYP*, and normalised to uninfested control plants. Fold-change, 2-AACt, is relative to the expression of cv. Concerto at 0 h. Boxplots show median and confidence intervals; total number of model observations for each gene = 20.

**Table 3:**
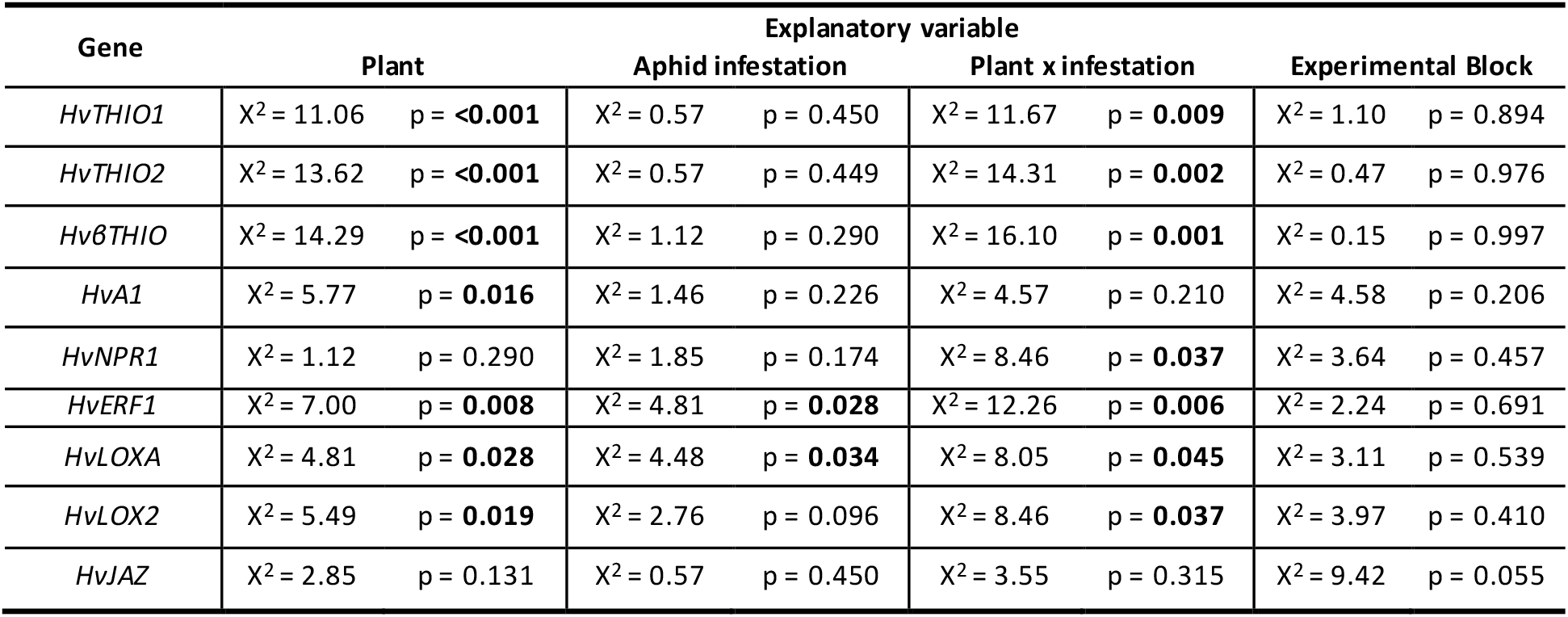
Statistical results of plant defence gene expression assessed by qPCR showing χ^2^ and p values (significant values are shown in bold)

No genes were differentially expressed in response to aphid infestation. However, several of the phytohormone signalling genes were differentially expressed in response to the plant x aphid infestation interaction; namely *HvLOXA, HvLOX2, HvNPR1*, and *HvERF1* (Table 3). *HvLOXA* was significantly down-regulated in Hsp5 after 24h aphid infestation to levels similar to those observed in Concerto (Fig. 6; Table 4). For *HvLOX2, HvNPR1*, and *HvERF*, expression levels were higher in Hsp5 after 24h of aphid infestation compared with levels in Concerto (Fig. 6; Table 4). *HvJAZ* was not differentially regulated in response to any treatment factor.

**Table 4:**
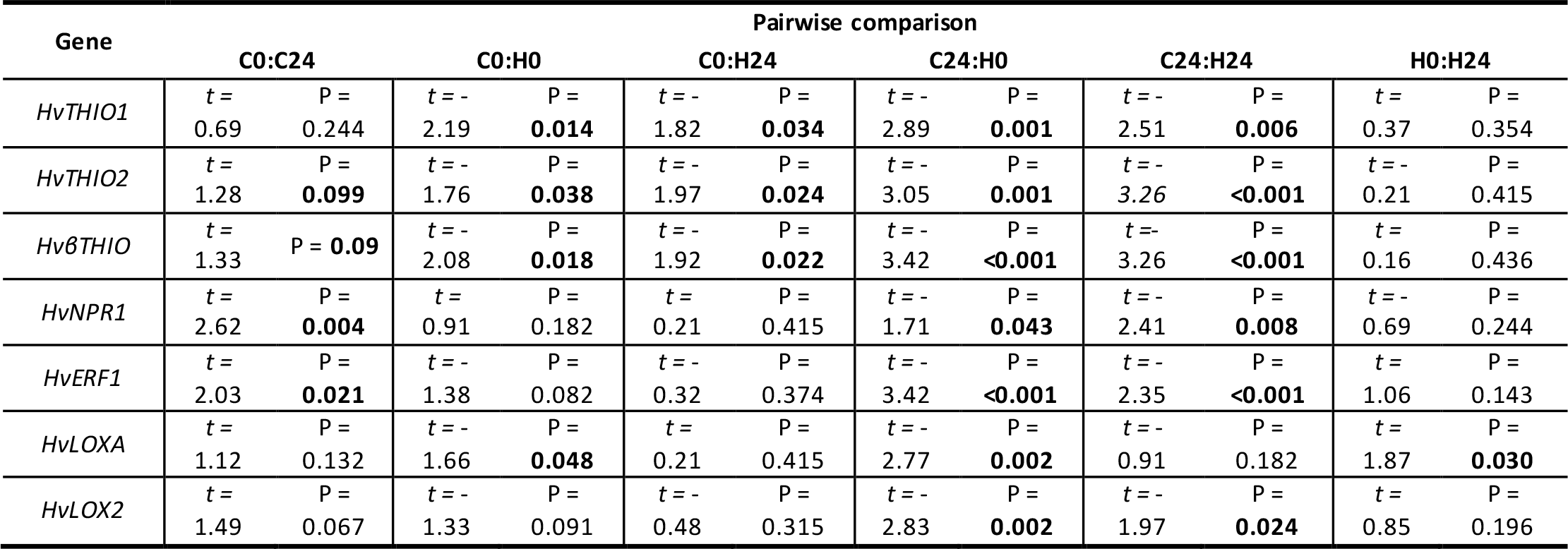
Dunn’s test *post-hoc* analysis of qPCR results significant for plant x time-point interaction, showing *t* and p values for each pairwise comparison; C = Concerto, H = Hsp5, 0 = 0 h infestation, 24 = 24 h aphid infestation (significant values are shown in bold).

### Phloem amino acid composition in Hsp5 is characterised by a reduction in the proportion of essential amino acids and an increased abundance of asparagine

To investigate phloem factors contributing to partial-resistance against aphids (Fig. 4; Table 2) we analysed the amino acid composition and the proportion of essential and non-essential amino acids in phloem exudates (Fig. 7). Principal component analysis of amino acid composition (mol% of Asp, Glu, Asn, His, Ser, Gln, Arg, Gly, Thr, Tyr, Ala, Trp, Met, Val, Phe, Ile, Leu, and Lys) of phloem exudates revealed that three principal components explained 69% of the observed variation in overall amino acid composition, with all three principle components showing a degree of separation between the two plant types (Table 5). Furthermore, >50% of the variation was explained by the first two principal components (Fig. 7C; Table 5) with separation of Hsp5 from Concerto largely occurring along PC1 due to differences in the relative amounts of Asn and His vs. most other essential amino acids (Fig. 7C). The total percentage of essential amino acids (Arg, His, Ile, Leu, Lys, Met, Phe, Thr, Trp, Val; described in Morris (1991)) was higher in Concerto compared with Hsp5 (18.97% vs 11. 45%; 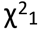 = 25.43; p = <0.001; Fig. 7D; Table 6).

The difference in composition of non-essential amino acids was most pronounced between the two plant types, with a higher proportion of Asn and lower proportions of Glu and Gly in Hsp5 (42.75%, 7.34% and 12.69%, respectively) compared with Concerto (7.55%, 15.18% and 22.43%, respectively) (Fig 7A-B). Amino acid composition and percentage of essential amino acids changed over the 24h of the experiment, but not in response to aphid treatment (Table 5; Table 6). Harvesting of leaf material for qPCR analysis had no effect on the phloem amino acid composition (Table 5; Table 6).

**Fig. 7:**
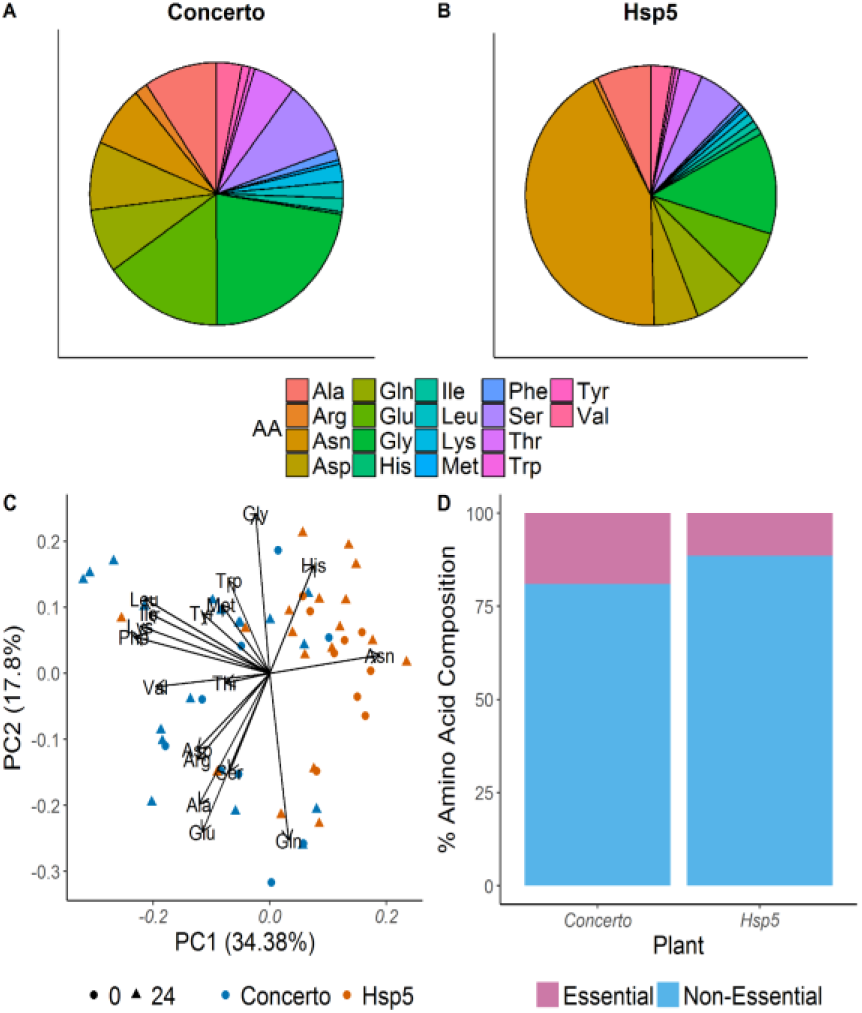
Amino acid composition of leaf phloem exudates of Hsp5 and Concerto. A and B) phloem amino acid composition in uninfested leaves of Concerto and Hsp5, respectively, at time zero. C) Biplot of scores on the first two principal components explaining >50% of the detected variation in amino acid composition. D) % proportion of essential and nonessential amino acids in phloem exudates of uninfested Concerto and Hsp5leaves at time zero.

**Table 5:** Analysis of deviance table for the scores on each principal component (% variation explained is indicated in brackets) derived from Principal Component Analysis of phloem amino acid composition (significant values are shown in bold)

| Explanatory Variable | PC1 (34.38%) |  | PC2 (17.80%) |  | PC3 (10.89%) |  |
| --- | --- | --- | --- | --- | --- | --- |
|  | X<sup>2</sup> (df) | p-value | X<sup>2</sup> (df) | p-value | X<sup>2</sup> (df) | p-value |
| Plant | 40.11 (1) | **<0.001** | 5.40 (1) | **0.020** | 11.98 (1) | **<0.001** |
| Aphid infestation | 0.29 (1) | 0.585 | 0.24 (1) | 0.626 | 2.26 (1) | 0.131 |
| Time-point | 5.42 (1) | **0.019** | 3.05 (1) | 0.080 | 0.08 (1) | 0.765 |
| Plant x time-point | 0.05 (1) | 0.816 | 2.17 (1) | 0.140 | 1.97 (1) | 0.159 |
| RNA Harvest | 0.17 (1) | 0.671 | 1.92 (1) | 0.166 | 0.67 (1) | 0.413 |

**Table 6:** Analysis of deviance table for essential amino acids as a percentage of total amino acids in phloem sap of Hsp5 and Concerto leaves (significant values are shown in bold).

| Explanatory Variable | Essential Amino acid (%) |  |
| --- | --- | --- |
|  | X<sup>2</sup> (df) | p-value |
| Plant | 23.35 (1) | **<0.001** |
| Aphid infestation | 0.13 (1) | 0.716 |
| Time-point | 8.82 (1) | **0.002** |
| Plant x time-point | 0.39 (1) | 0.530 |
| RNA Harvest | 2.01 (1) | 0.155 |

## Discussion

In this study we provide evidence for mesophyll and phloem mediated partial-resistance against aphids in a wild barley relative, which is effective against three aphid species, including two agricultural pests and an invasive species. Our data suggest the partial-resistance in this progenitor of modern barley is complex and may be associated with elevated defence gene expression and reduced phloem quality.

### Mesophyll and phloem-based factors contribute to partial-resistance in Hsp5

Our observation that *R. padi* feeding on Hsp5 spends more time in the pathway phase (probing into plant tissue) and takes longer to reach the phloem and initiate salivation/phloem ingestion is representative of feeding patterns associated with mesophyll-based resistance (Alvarez et al., 2006). Factors associated with mesophyll resistance are thought to antagonise and impede stylet penetration (Montllor and Tjallingii, 1989; Alvarez et al., 2006). In addition, *R. padi* spends less time ingesting phloem sap and salivating into the phloem, in line with resistance factors residing in the phloem (Alvarez et al., 2006). Phloem based resistance factors generally alter the ingestion efficiency of aphids and can involve feeding antagonists, alterations to the phloem metabolic profile, and the plugging of sieve-tube elements (Klingler et al., 1998; Klingler et al., 2005; Pegadaraju et al., 2007; Medina-Ortega and Walker, 2015; Greenslade et al., 2016; Peng and Walker, 2018).

Partial-resistance against aphids in other plant-aphid systems have similarly pointed to the importance of mesophyll and phloem-based resistance factors (Montllor and Tjallingii, 1989; Caillaud et al., 1995; Pegadaraju et al., 2007; Guo et al., 2012; Greenslade et al., 2016). For example, partial resistance in *Triticum monococcum* lines (a wild relative of wheat) against *S. avenae* and *R. padi* involved resistance at the phloem level, as determined by EPG (Caillaud et al., 1995; Greenslade et al., 2016). Interestingly, metabolic profiling of resistant and susceptible lines in the latter study indicated different metabolite profiles between susceptible and partially-resistant plants, including lower levels of primary metabolites in the partially-resistant lines and elevated Asn (Greenslade et al., 2016). Resistance against *D. noxia* in five cereal species was shown to involve phloem and mesophyll resistance factors, with elevated 2,4-dihydroxy-7-methoxy-1,4-benzoxazin-3-one (DIMBOA) levels hypothesised to be a contributing factor towards resistance (Mayoral et al., 1996). Resistance against *D. noxia* in wheat also involved individual resistance genes; the most recent of these to be characterised is *Dn10* (Li et al., 2018), although the tissue types involved in conferring this resistance are unknown.

In addition, *Arabidopsis thaliana* defence against *Myzus persicae* involved *phytoalexin deficient4* (*PAD4*), and reduced susceptibility in *PAD4* overexpressing lines was associated with reduced phloem ingestion (Pegadaraju et al., 2007). Furthermore, gene-for-gene type resistance against the pea aphid, *Acyrthosiphonpisum*, in *Medicago truncatula* line Jemalong A17 (A17) is conferred by a single gene, *RAP1*, which maps to chromosome 3 (Stewart et al., 2009). Further characterisation showed that A17 also conferred resistance against the bluegreen aphid, *A. kondoi*, as well as *A. pisum*, and that this resistance was phloem-mediated and linked to a QTL on chromosome 3 (Guo et al., 2012). The inorganic chemistry of leaf tissue can also play a role in plant defence (Lane, 2002; Boyd, 2012), with oxalates highlighted as important contributors towards plant defence against pathogens in cereals (Lane, 2002). The inorganic chemistry of Hsp5 and Concerto leaves was not assessed in this study, but a comparison of leaf chemistry could highlight further chemical groups which could be involved in mediating plant-aphid interactions.

### Differences in leaf surface characteristics contribute little to partial-resistance against aphids in Hsp5

Leaf surface architectural differences were detected between Concerto and Hsp5 and included a higher abundance of non-glandular leaf trichomes in Hsp5 and a more complex chemistry in the epicuticular extracts of Concerto. Non-glandular trichomes have been reported to decrease *R. padi* fitness in wheat (Roberts and Foster, 1983), but have contrasting effects on herbivore fitness in other plants. For example, the density of non-glandular trichomes in raspberry negatively affected spider mite performance but had minimal impact on aphid fitness (Karley et al., 2016).

Although we detected a positive correlation between trichome abundance and *R. padi* rm, there was no correlation between trichome abundance and juvenile *R. padi* mass gain, indicating that either non-glandular trichomes have differential effects on aphids at different life-stages or that the higher abundance of non-glandular trichomes on the leaf surface of Hsp5 are not a primary cause of partial-resistance to aphids. Indeed, EPG analysis indicated that leaf surface traits contributed little to partial-resistance in Hsp5. Aphids feeding from Hsp5 did not show increased time spent in the non-probing phase (a key indicator of epidermal-mediated resistance: Alvarez et al. (2006). Moreover, the time taken for aphids to penetrate the leaf surface was shorter on Hsp5, indicating aphids experienced fewer barriers to probing the leaf epidermis on Hsp5. Less complex chemistry at the leaf epicuticle in Hsp5 may have played a role in promoting aphid probing of plant tissue and previous studies have shown that epicuticular chemical compounds can either promote or deter aphid probing of plant tissue (Powell et al., 1999). Barley epicuticular waxes can contribute towards resistance against *R. padi* (Tsumuki et al., 1989), although the influence of specific chemical functional groups on aphid behaviour is not well characterised.

### Basal defence gene expression is elevated in Hsp5

In line with Delp et al. (2009) and Mehrabi et al. (2014), basal expression levels of thionin genes, which contribute to barley defences against aphids (Escudero-Martinez et al., 2017), and *LOX2*, a JA-signalling marker, were elevated in Hsp5. The feeding parameters exhibited by *R. padi* while feeding on Hsp5 indicated that mesophyll-based mechanisms conferred short-term (6h) partial-resistance against aphids. Thionins are antimicrobial peptides present in plant cell walls and within the intracellular space (Reimann-Philipp et al., 1989; Stec, 2006). It is therefore likely that aphids are exposed to thionins when probing mesophyll tissue, and these could contribute to reduced aphid performance on Hsp5.

In addition, several plant defence signalling genes, *LOX* (JA), *ERF1* (ET) and *A1* (ABA) showed higher basal expression in Hsp5, suggesting that Hsp5 may be able to respond more efficiently to biotic stimuli. In line with our data, *HvLOX2* genes and other components of the JA (*HvAOS, HvAOC*), ET (*HvACCO, HvACCS*), and Auxin (*HvTDS, HvTSM*) signalling pathways showed higher basal expression in the barley cultivar Stoneham, which is partially-resistant against *D. noxia*, relative to susceptible cultivar Otis (Marimuthu and Smith, 2012). Furthermore, overexpression of *HvLOX2* in barley resulted in decreased *R. padi* fecundity five days after infestation (Losvik et al., 2017), pointing to a functional role of this gene and JA signalling in short-term barley defence against aphids. Expression levels of *CmERF1* in early responses (6h) to aphid infestation were around ten-fold higher in a resistant variety of melon compared with a susceptible variety (Anstead et al., 2010), and higher ET levels have been reported to contribute to *R. maidis* resistance in maize (Louis et al., 2015). ABA mediated-processes have been implicated in increasing both plant resistance (Zhu-Salzman et al., 2004; Park et al., 2006) and susceptibility (Kerchev et al., 2013; Hillwig et al., 2016) to aphids. Therefore the consequence of elevated *HvA1* in the context of partial aphid resistance in Hsp5 is not clear. Guo et al. (2016) reported that elevated ABA levels in response to drought conditions can lead to increased aphid xylem ingestion. This finding suggests that elevated expression of components of the ABA signalling pathway could indirectly contribute to partial-resistance by encouraging xylem ingestion, thereby reducing phloem ingestion. Indeed, longer periods of xylem ingestion were detected in *R. padi* feeding from Hsp5, indicating that there may be potential fitness consequences for aphids as a result of higher basal expression levels.

In contrast, basal expression levels of *HvNPR1*, a SA-signalling marker, were not significantly different between Hsp5 and Concerto. However, *HvNPR1* levels in Concerto were down-regulated 24h after aphid infestation to a level that was significantly lower than *HvNPR1* levels in Hsp5 after 24h of aphid infestation. Therefore, SA-mediated defences in Concerto, but not Hsp5, may be repressed upon aphid infestation leading to increased susceptibility of this cultivar. Indeed, NPR1 is required for initiation of SA-mediated defence against aphids in *A. thaliana* (Moran and Thompson, 2001; Wu et al., 2012).

### Essential amino acid availability is reduced in Hsp5

The observation that the relative concentration of essential amino acids was lower in the phloem of Hsp5 compared with Concerto could be associated with lower aphid performance.

Alterations to amino acid composition can reduce plant palatability and nutritional quality and contribute to increased resistance against aphids (Sandström and Pettersson, 1994; Ponder et al., 2000; Karley et al., 2002). For example, decreased survival and fecundity of *M. persicae* and *Macrosiphum euphorbiae* feeding from older “tuber-filling” potato plants compared with younger “pre-tuber-filling” plants was linked to changes in amino acid composition (Karley et al., 2002). When presented with chemically-defined diets representative of the phloem sap composition of young and mature potato plants, *M. persicae* and *Ma. euphorbiae* showed decreased feeding rate on the “tuber-filling” diets (Karley et al., 2002). In addition, *R. padi* feeding from barley plants grown under nitrogen-limited conditions exhibited reduced rm compared with *R. padi* feeding on barley grown under nitrogen-rich conditions (Ponder et al., 2000). Examination of phloem amino acid composition indicated that composition was altered under nitrogen-limited conditions. Interestingly, plants under nitrogen-limited conditions contained a higher percentage of Asn, a lower percentage of Gly and a small reduction in the concentration of essential amino acids compared with nitrogen-fertilised plants (Ponder et al., 2000).

Changes in the abundance of specific amino acids cannot be readily associated with increased insect resistance/susceptibility in a given plant species. It is most likely that any observed effect on aphid fitness is due primarily to the ratio of essential:non-essential amino acids as a result of compositional changes. Indeed, essential amino acids are generally elevated in susceptible plants compared with varieties showing increased resistance against aphids (Auclair, 1976). Vogel and Moran (2011) found that the mass of multiple *A. pisum* biotypes was reduced when essential amino acids were removed from aphid artificial diets; this was observed even though aphids have the capacity to synthesis essential amino acids via their essential endosymbiont, *Buchnera aphidicola* (Douglas and Prosser, 1992). Furthermore, some aphid species actively remobilise plant nutrients to increase the abundance of essential amino acids in susceptible plant hosts (Telang et al., 1999; Sandström et al., 2000). Enhanced mobilisation of plant nutrients is thought to be the mechanism explaining resistance breakdown in wheat by a virulent biotype of the greenbug, *Schizaphis graminum* (Dorschner et al., 1987). In a study by Sandström et al. (2000), *R. padi* feeding did not lead to increased phloem mobilisation of amino acids, similar to observations made in our experimental system where *R. padi* infestation did not alter the phloem amino acid composition in Hsp5 or Concerto.

## Conclusion

This study shows that partial-resistance against aphids in wild barley can be wide-ranging and effective against multiple aphid species. Broad partial-resistance traits are agriculturally and economically important as crops are likely to be infested by multiple insect species under field conditions (Stam et al., 2014). However, compared with gene-for-gene resistance traits, comparatively little is known about the genetic determinants of partial-resistance. The underlying factors contributing towards partial-resistance are complex and likely to involve multiple, independent resistance pathways (Mayoral et al., 1996; Moharramipour et al., 1997; Seah et al., 1998). We show that partial-resistance against aphids in wild barley involves resistance factors that reside in both the mesophyll and vascular tissues. This partial-resistance is potentially caused by heightened basal expression of defence and phytohormone signalling genes, alongside altered phloem amino acid quality. However, the functional molecular role of these differences, and how they contribute towards partial-resistance, remains to be established.

## Supplementary Data

Supplementary Table 1: primers used in RT-qPCR analysis.

Supplementary Table 2: Additional EPG parameters.

Supplementary Table 3: FTIR results for leaf surface extracts

Supplementary Fig. 1: Correlation of insect fitness with non-glandular trichome density.

## Acknowledgements

D.J.L was funded by the James Hutton Institute and the Universities of Aberdeen and Dundee through a Scottish Food Security Alliance (Crops) PhD studentship. A.J.K, T.A.V, A.H.J.R, A.M.M, and E.P-F were supported by the strategic research programme funded by the

Scottish Government’s Rural and Environment Science and Analytical Services Division. J.I.B.B. was supported by the European Research Council (310190-APHIDHQST). We would like to thank Dr Freddy Tjallingii (EPG Systems, The Netherlands), Professor Alberto Fereres (CSIC, Spain) and Professor Gregory Walker (University of California, Riverside, USA) for providing EPG training and Gaynor Malloch (James Hutton Institute) for kindly providing COI barcoding primers.

## Author Contributions

JIBB, AJK, and DJL conceived and designed the experiments. DJL, EP-F, and AMM performed the experiments. JIBB, AJK, DJL, and JAHR analysed the data. DJL wrote the manuscript with input from JIBB, AJK, TAV, and JAHR. All authors read and approved the final manuscript.

## Abbreviations

ABA: Abscisic Acid
Ala: Alanine
Arg: Arginine
Asn: Asparagine
Asp: Aspartic Acid
Concerto: *Hordeum vulgare* cv.Concerto
EPG: Electrical penetration graph
ET: Ethylene
FTIR: Fourier Transform Infrared
Gln: Glutamine
Glu: Glutamic Acid
Gly: Glycine
His: Histidine
Hsp5: *Hordeum spontaneum*5
Ile: Isoleucine
JA: Jasmonic Acid Leu: Leucine
Lys: Lysine
Met: Methionine
Phe: Phenylalanine
r_m_: the intrinsic rate of population increase
SA: Salicylic Acid
Ser: Serine
Thr: Threonine
Trp: Tryptophan
Tyr: Tyrosine
Val: Valine

## Notes

#### Summary of Updates

Inclusion of Supplementary Material

