## Supplementary Table 1, Supplementary Table 2, Supplementary Table 3, Supplementary Fig. 1 for "Partial-resistance against aphids in wild barley involves phloem and mesophyll-based defences"

Supplementary table 1: primers used in qPCR analysis

| Gene | Name | Accession no. | Forward primer | Reverse Primer | Primer Reference |
|---|---|---|---|---|---|
| *HvUBC* | Ubiquitin-conjugating enzyme | AK248472.1 | TCAATTCCCGAGCAGTATCC | AGATTGCCTGAGTCGCAGTT | Hua et al 2014 |
| *HvCYP* | Cytochrome P450 | AK253120.1 | CTGTCGTGTCGTCGGTCTAA | TGAAAGCGACAAACAGATGC | Hua et al 2014 |
| *HvLOXA* | Lipoxygenase 1 | AY220737.2 | gccagatccagaccatcatc | tcggaggagtgcttcgac | Designed for this study |
| *HvLOX2* | Predicted Lipoxygenase | AK357253.1 | atgtcctatcccacgacacc | agtgcgtcctcagccagt | Escudero-Martinez et al 2017 |
| *HvJAZ* | Jasmonate ZIM-domain protein 3 | MLOC_9995.2 | aggaaaagtggtcgtggttg | atctggagcaatccgttgac | Escudero-Martinez et al 2017 |
| *HvA1* | ABA-inducible late embryogenesis abundant protein | X13498.1 | atgggaggggacaacacc | ggaaattaagcgcgaacg | Designed for this study |
| *HvNPR1* | Non-expresser of pathogenesis-related genes 1-Like | MLOC_64922.1 | ttgataacatctagaggcaatgct | tgcgtgaaactgttcgagag | Designed for this study |
| *HvERF1* | Ethylene-response factor 1 | HQ328941.1 | ctatataatgattgggtgcatgttg | ggcatatgacccaaggtgtt | Designed for this study |
| *HvTHIO1* | Thionin 1 | AK359149 | tatggccaaggtcgttttgt | cataactaagatgatacatttgcttcg | Escudero-Martinez et al 2017 |
| *HvTHIO2* | Thionin 2 | AK357884 | gcggttcaaaatgtcctagtg | ccaatggtgcagtactgagtg | Escudero-Martinez et al 2017 |
| *HvβTHIO* | β Purothionin | AK252675.1 | tactgggtttagttctggagcag | acgtgtccttgcagcaactt | Escudero-Martinez et al 2017 |

Supplementary table 2: Additional EPG parameters

| EPG Parameter | Hypothesised location of resistance factor | Transformation | Mean Value Concerto | Mean Value Hsp5 | Statistical Test | Test Statistic | p value |
|---|---|---|---|---|---|---|---|
| average non probing (period duration) | Epidermis | - | 804.11 s | 442.26 s | K.W | 0.87 | 0.352 |
| median non probing (period duration) | Epidermis | - | 654.92 s | 380.85 s | K.W | 0.80 | 0.352 |
| sum of non probing | Epidermis | sqrt | 2552.92 s | 2095.83 s | ANOVA | 0.51 | 0.483 |
| number of non probing periods | Epidermis | log[2] | 5.22 | 0.89 | GLM | 0.40 | 0.534 |
| number of brief probes < 3 min before 1st E | Epidermis, Mesophyll | - | 0.11 | 0.81 | K.W | 2.17 | 0.141 |
| number of probes before 1st pd | Epidermis, Mesophyll | - | 0.66 | 0.87 | K.W | 1.98 | 0.159 |
| number of probes before the first G | Epidermis, Mesophyll | - | 1.94 | 1.87 | K.W | 0.50 | 0.480 |
| number of probes | Epidermis, Mesophyll | log[2] | 5.22 | 5.62 | GLM | 0.40 | 0.534 |
| sum of probing | Epidermis, Mesophyll | - | 19057.08 s | 19504.17 s | K.W | 0.43 | 0.558 |
| sum of C | Epidermis, Mesophyll | sqrt | 7357.38 s | 8051.17 s | ANOVA | 0.30 | 0.590 |
| median time to 1st pd in all probes with a pd | Epidermis, Mesophyll | sqrt | 112.47 s | 118.31 s | ANOVA | 0.28 | 0.605 |
| number of brief probes (probes < 180 s) | Epidermis, Mesophyll | - | 0.61 | 1.06 | K.W | 0.80 | 0.949 |
| number of C periods | Epidermis, Mesophyll | - | 11.00 | 11.12 | GLM | 0.01 | 0.950 |
| median probe | Epidermis, Mesophyll | - | 5691.88 s | 3627.73 s | K.W | 0.01 | 0.973 |
| min. time to 1st pd in 1st probe | Epidermis, Mesophyll | - | 60.47 s | 60.53 s | K.W | 0.00 | 0.973 |
| no. pd per min C , only C_phases with pd | Mesophyll | - | 0.54 | 0.67 | GLM | 1.85 | 0.183 |
| sum of pd | Mesophyll | sqrt | 353.79 | 416.04 | ANOVA | 1.54 | 0.223 |
| duration of the first pd | Mesophyll | - | 4.27 s | 5.33 s | K.W | 1.46 | 0.227 |
| no. pd per min C | Mesophyll | - | 0.65 | 0.75 | GLM | 0.97 | 0.332 |
| median duration of pd | Mesophyll | - | 4.05 s | 4.72 s | K.W | 0.93 | 0.334 |
| number of F | Mesophyll | - | 0.17 | 0.06 | K.W | 0.86 | 0.354 |
| time to 1st pd (from start of 1st probe) | Mesophyll | sqrt | 220.34 s | 289.85 s | ANOVA | 0.80 | 0.378 |
| mean duration of the first 5 pd | Mesophyll | - | 4.30 s | 5.07 s | K.W | 0.74 | 0.388 |
| average F | Mesophyll | - | 492.66 s | 825.67 s | K.W | 0.69 | 0.405 |
| median F | Mesophyll | - | 492.66 s | 825.67 s | K.W | 0.69 | 0.405 |
| sum of F | Mesophyll | - | 492.66 s | 825.67 s | K.W | 0.69 | 0.405 |
| average duration of pd | Mesophyll | - | 4.34 s | 4.97 s | K.W | 0.68 | 0.408 |
| number of pd | Mesophyll | - | 72.61 | 83.12 | K.W | 0.58 | 0.448 |
| median C | Mesophyll | log[2] | 502.91 s | 459.90 s | ANOVA | 0.25 | 0.618 |
| average probe | Mesophyll | log[2] | 6802.31 s | 4884.68 s | ANOVA | 0.25 | 0.622 |
| duration of the second pd | Mesophyll | - | 4.45 s | 5.29 s | K.W | 0.22 | 0.641 |
| average C; with pd without E1e, F and G | Mesophyll | - | 742.72 s | 763.48 s | ANOVA | 0.03 | 0.866 |
| time to 1st pd in 1st probe with a pd | Mesophyll | sqrt | 205.05 s | 160.95 s | ANOVA | 0.01 | 0.894 |
| average time to 1st pd in all probes with a pd | Mesophyll | sqrt | 153.11 s | 131.63 s | ANOVA | 0.01 | 0.924 |
| time to 1st E within the 1st probe with E | Mesophyll, Phloem | log[10] | 1018.99 s | 1453.15 s | ANOVA | 1.35 | 0.253 |
| number of probes before 1st sE2 | Mesophyll, Phloem | - | 1.72 | 1.68 | K.W | 0.59 | 0.442 |
| number of probes after 1st sE2 | Mesophyll, Phloem | - | 2.17 | 1362.00 | K.W | 0.59 | 0.442 |
| time from the 1st E1 to 1st E2 | Mesophyll, Phloem | - | 1538.73 s | 2493.86 s | K.W | 0.39 | 0.535 |
| minimum time to 1st E within probes | Mesophyll, Phloem | - | 687.47 s | 1043.44 s | K.W | 0.17 | 0.678 |
| average time to 1st E within probes | Mesophyll, Phloem | - | 912.91 s | 1581.33 s | K.W | 0.12 | 0.729 |
| number of all E1 periods (sgE1 + frE1) | Mesophyll, Phloem | log[2] | 5.44 | 4.56 | GLM | 0.11 | 0.749 |
| Total time in E1 before E2 | Phloem | - | 825.60 s | 300.07 s | K.W | 3.37 | 0.066 |
| E2 index: % | Phloem | - | 0.50 | 0.35 | K.W | 3.11 | 0.078 |
| Total time in E1 | Phloem | log[10] | 1029.75 s | 459.12 s | ANOVA | 3.08 | 0.089 |
| sum of fractions of E1 | Phloem | - | 858.83 | 320.75 | K.W | 2.86 | 0.091 |
| number of E12 phloem periods i.e. with both, E1 and E2 | Phloem | sqrt | 2.88 | 1.93 | GLM | 3.03 | 0.091 |
| median E2 | Phloem | - | 4803.10 | 1601.25 | K.W | 2.52 | 0.113 |
| number of E2 periods | Phloem | sqrt | 2.88 | 2.00 | GLM | 2.62 | 0.115 |
| median E12 (with both E1 and/or E2) | Phloem | - | 5363.61 s | 1751.12 s | K.W | 2.30 | 0.129 |
| number of fractions of E1;: E1followed/preceded by E2 | Phloem | sqrt | 2.88 | 2.06 | GLM | 2.26 | 0.142 |
| maximum E1 period (either sgE1 or frE1) | Phloem | - | 762.95 s | 225.69 s | K.W | 2.10 | 0.147 |
| maximum duration of a fraction of E1 | Phloem | - | 738.92 s | 215.04 s | K.W | 2.00 | 0.157 |
| average E1 | Phloem | - | 334.70 s | 87.26 s | K.W | 2.00 | 0.157 |
| median E1 (sgE1 and E1fr) | Phloem | log[10] | 290.67 s | 69.61 s | ANOVA | 1.70 | 0.202 |
| E1 index:duration E1/ allE as % | Phloem | - | 0.16 | 0.25 | K.W | 1.46 | 0.227 |
| average fraction of E1 | Phloem | - | 593.51 | 117.22 | K.W | 0.69 | 0.408 |
| median fraction of E1 | Phloem | - | 550.17 | 95.72 | K.W | 0.53 | 0.469 |
| median single E1 | Phloem | sqrt | 47.17 s | 47.32 s | ANOVA | 0.44 | 0.513 |
| number of single E1 (without E2) periods | Phloem | sqrt | 2.55 | 2.50 | GLM | 0.37 | 0.545 |
| average single E1 | Phloem | sqrt | 52.66 s | 49.33 s | ANOVA | 0.20 | 0.658 |
| number of E2 before the 1st sE2 | Phloem | - | 0.72 | 0.56 | K.W | 0.16 | 0.685 |
| sum of E2 before 1st sE2 | Phloem | - | 123.11 s | 127.56 s | K.W | 0.09 | 0.762 |
| maximum duration of a single E1 period | Phloem | sqrt | 85.07 s | 75.23 s | ANOVA | 0.08 | 0.774 |
| sum of single E1 | Phloem | - | 170.92 s | 138.37 s | K.W | 0.05 | 0.809 |
| mean duration of E2 periods before the 1st sE2 | Phloem | - | 59.25 s | 65.46 s | K.W | 0.02 | 0.900 |
| time to the first G (after first penetration) | Mesophyll, Xylem | - | 10761.03 s | 6577.95 s | K.W | 1.22 | 0.270 |

| | | | | | | | |
|---|---|---|---|---|---|---|---|
| Average xylem ingestion time | Xylem | - | 2117.75 s | 4043.78 s | K.W | 3.03 | 0.082 |
| median G | Xylem | - | 2117.75 s | 3770.59 s | K.W | 2.24 | 0.134 |

Supplementary table 3: Functional groups identified using FTIR spectral analysis on dichloromethane leaf surface extracts for Hsp5 and Concerto, showing presence of band ($cm^{-1}$) in the spectra and allocated functional group

| **Plant** | **Rep** | **Number of functional groups** | **Identified Bands ($cm^{-1}$) and Functional Groups** | | | | | | | | |
|---|---|---|---|---|---|---|---|---|---|---|---|
| HsP5 | 1 | 3 | 2959 $CH_3$ stretching | 2920/2852 $CH_2$ stretching | 1736 C=O stretch for ester group (v. weak) | | | | | 1473/1462 $CH_2$ deformation | 731/719 $CH_2$ wag |
| | 2 | 2 | 2959 $CH_3$ stretching | 2920/2852 $CH_2$ stretching | | | | | | 1473/1462 $CH_2$ deformation | 731/719 $CH_2$ wag |
| | 3 | 2 | 2959 $CH_3$ stretching | 2920/2852 $CH_2$ stretching | | | | | | 1473/1462 $CH_2$ deformation | 731/719 $CH_2$ wag |
| | 4 | 3 | 2959 $CH_3$ stretching | 2920/2852 $CH_2$ stretching | 1736 C=O stretch for ester group (v. weak) | | | | | 1473/1462 $CH_2$ deformation | 731/719 $CH_2$ wag |
| Concerto | 1 | 4 | 2959 $CH_3$ stretching | 2920/2852 $CH_2$ stretching | 1736 C=O stretch for ester group | | | 1578/1540 $RCOO^-$ carboxylic acid salt | | 1473/1462 $CH_2$ deformation | 731/719 $CH_2$ wag |
| | 2 | 5 | 2959 $CH_3$ stretching | 2920/2852 $CH_2$ stretching | 1736 C=O stretch for ester group | | 1605/1515 C=C stretch aromatic compound | 1581/1390/ 1263 $RCOO^-$ carboxylate | | 1473/1462 $CH_2$ deformation | 731/719 $CH_2$ wag |
| | 3 | 5 | 2959 $CH_3$ stretching | 2920/2852 $CH_2$ stretching | 1736 C=O stretch for ester group | 1653/1549 amide I and II protein or amide | | | 1354 nitro containing component (possibly $NO_3$) | 1473/1462 $CH_2$ deformation | 731/719 $CH_2$ wag |
| | 4 | 3 | 2959 $CH_3$ stretching | 2920/2852 $CH_2$ stretching | 1736 C=O stretch for ester group | | | | | 1473/1462 $CH_2$ deformation | 731/719 $CH_2$ wag |

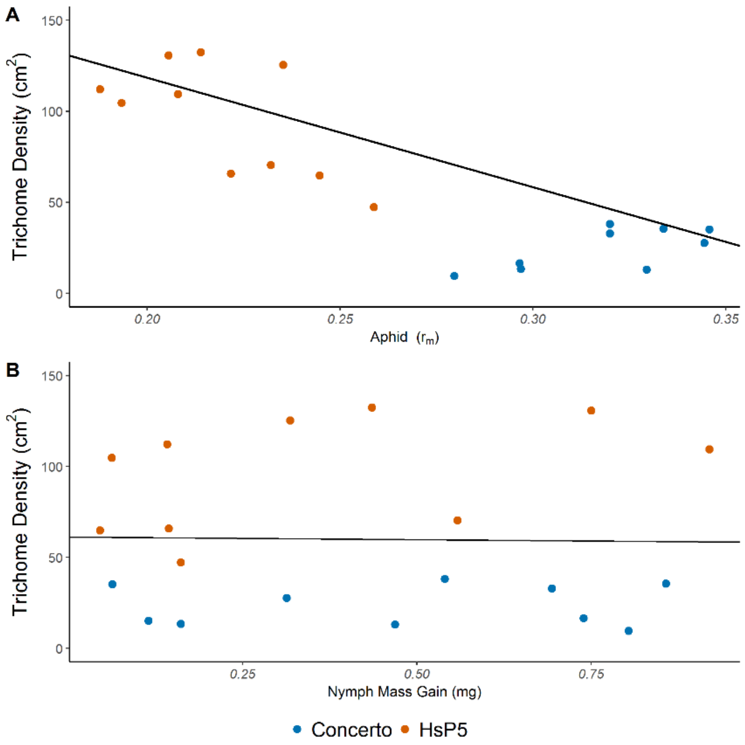


Supplementary Fig. 1: Comparison of insect fitness with plant trichomes. A) Correlation between leaf non-glandular trichome density ($cm^2$) and *R. padi* $r_m$. Line represents model correlation coefficient. Number of model observations = 19. B) Correlation between leaf non-glandular trichome density ($cm^2$) and *R. padi* nymph mass gain. Line represents model correlation coefficient. Number of model observations = 20
